# Accelerated Limbal Epithelial Differentiation of Human Induced Pluripotent Stem Cells Using a Defined Keratinocyte Serum-Free Medium

**DOI:** 10.1101/2024.08.06.606916

**Authors:** Danial Roshandel, Belén Alvarez-Palomo, Khine Zaw, Dana Zhang, Michael J Edel, Evan N Wong, Steven Wiffen, Fred K Chen, William Morgan, Samuel McLenachan

## Abstract

**Purpose:** Treatment of bilateral limbal stem cell deficiency (LSCD) is challenging due to the limited autologous stem cell sources. This study aimed to differentiate human induced pluripotent stem cells (hiPSCs) into limbal epithelial stem cells (LESCs) using a defined keratinocyte serum-free medium (DKSFM).

**Methods:** A fully characterized hiPSC line was committed to ectodermal differentiation using Essential 6 (E6) medium supplemented with 10 µM Y-27632 (Day 1), 10 µM SB-505124 plus 50 ng/ml bFGF (Day 2) and 25 ng/ml BMP-4 (Days 3 and 4). Differentiation was continued in DKSFM for an additional 21 days. Quantitative PCR (qPCR) and/or immunocytochemistry (ICC) for pluripotency, proliferation, LESC, and corneal epithelial markers were performed on samples collected at days 5, 10, 15, and 25 (D5 to D25) and compared with undifferentiated hiPSCs (UD).

**Results:** qPCR revealed a significant decrease in the expression of *OCT4* and *NANOG* and a significant increase in *ABCG2* and *TP63* following ectodermal induction (i.e., D5), compared with UD (P < 0.05). The expression levels of *Ki67*, *ABCG2*, *TP63*, and *CK14* were significantly higher at D10, compared with D5 and D25 (P < 0.05). The ratio of p63α-positive cells was 71% and 56% in D10 and D15 cells, respectively (P < 0.05).

**Discussion:** Our method resulted in a limited but rapid differentiation of hiPSCs into LESC-like cells. The LESC-like cells appeared as early as 5 days following ectodermal induction and their population peaked after 10 days. Upon further optimization and validation, DKSFM can be used for rapid limbal epithelial differentiation of hiPSCs.

## Introduction

Limbal epithelial stem cells (LESCs) are responsible for the constant replenishment of the corneal epithelium via continuous self-renewal and differentiation into corneal epithelial cells (CECs), a process which plays a crucial role in maintaining the ocular integrity and optimal vision.^1^ Limbal stem cell deficiency (LSCD) is mostly caused by ocular surface burns, but can also result from genetic defects (e.g. *PAX6* mutations), prolonged soft contact lens wearing, chronic autoimmune/inflammatory ocular surface diseases, ocular surgeries, or topical medications.^2^ Total LSCD is characterized by corneal opacification, vascularization, and epithelial defects, endangering the eye with potential devastating complications such as corneal ulceration and perforation, and endophthalmitis.^3^ Limbal stem cell transplantation (LSCT) is the current standard management for total LSCD. The standard procedure for bilateral cases includes allogenic LCST, which is associated with a significant risk of allograft rejection despite high-dose systemic immunosuppression.^4^ In addition, immunosuppression carries a high risk of systemic side effects, and may not be tolerated by some patients.^5^

Autologous stem cells can produce cells with anatomical and functional properties like CECs, while eliminating the need for immunosuppression. Oral mucosal epithelial stem cells (OMESCs) have been successfully used for regenerating the ocular surface epithelium, though the phenotype of the resultant epithelium, and its long-term stability, transparency and visual outcomes are yet to be explored.^6^ Hence, research on potential autologous sources of stem cells is still ongoing. Mesenchymal stem cells (MSCs) have been studied extensively as a potential candidate for tissue regeneration. However, MSCs have a mesodermal origin and might be difficult to differentiate to ectodermal cells such as corneal epithelium.^7^ In addition, although embryonic stem cells (ESCs) have been successfully differentiated into various cells including limbal and corneal epithelial cells, they might be rejected by the recipient.^8^ In recent years, induced pluripotent stem cells (iPSCs) have been efficiently differentiated into various cells and have shown promising outcomes in treating refractory diseases such as alloimmune platelet transfusion refractoriness in a human trial.^9^ In the eye, iPSC-derived retinal pigment epithelial (RPE) cells are under investigation in phase I/II clinical trials for the treatment of advanced age-related macular degeneration (www.clinicaltrials.gov NCT04339764 and NCT05445063; accessed on 7 August 2023). However, corneal epithelial regeneration using iPSC derived LESCs (iLESC) was hindered by a lack of a standard differentiation protocol and *in vivo* data.

Different protocols for the differentiation of human iPSCs (hiPSCs) into LESCs have been developed by several groups. Unlike earlier reports that used animal or human-based products such as PA6 feeder layer and limbal fibroblast-conditioned medium^10,11^, more recent protocols have used feeder and serum-free media to achieve better repeatability and consistency and safety for future clinical use.^12^ Based on the robust evidence supporting the role of Wnt, transforming growth factor beta (TGF-β), and fibroblast growth factor (FGF) pathways on corneal epithelial differentiation, Mikhailova and colleagues developed a two-stage protocol for induction of ectodermal commitment and epithelial differentiation using small molecules followed by further corneal epithelial differentiation in a defined medium.^12^ The protocol was further optimized by adding bone morphogenic factor 4 (BMP-4) to the induction medium.^13^ After three weeks of differentiation, more than 60% of cells expressed LESC markers (p63α/ΔNp63α). More interestingly, up to 90% of cells were positive for LESC markers after thawing of the cryopreserved cells.^14^

In the current study, we adopted the induction protocol suggested by Hongisto et al.^14^ with some modifications and used a defined keratinocyte serum-free medium for differentiating hiPSCs into corneal epithelial-like cells for potential future *in vivo* applications.

## Materials and methods

### Human iPSC line

A fully characterized hiPSC line reprogrammed from skin fibroblasts of a healthy control subject was obtained from the Ocular Tissue Engineering Laboratory biobank (Lions Eye Institute, Perth, Australia). The study protocol was approved by the institutional review board and the Human Research Ethics Committee of the University of Western Australia (2021/ET000151) and Sir Charles Gairdner and Osborne Park Health Care Group (RGS0000005680) and adhered to the tenets of Helsinki. The process of sampling, fibroblast isolation, culture and iPSC generation has been published elsewhere.^15,16^ Briefly, dermal fibroblasts were isolated from the skin biopsy and cultured in Dulbecco’s Modified Eagle’s Medium (DMEM, ThermoFisher) supplemented with 10% fetal calf serum (FBS) and antibiotic–antimycotic (15240096, ThermoFisher). Reprogramming was performed using Episomal iPSC Reprogramming Plasmid Kit (SC900A-1, System Biosciences), according to the manufacturer’s instructions and the iPSCs were characterized using pluripotency markers as described^16^ and stored in liquid nitrogen.

### Human corneal epithelial cell line

A human corneal epithelial cell (HCEC) line was obtained from ATCC (PCS-700-010, VA, USA) and cultured in the recommended medium (Corneal Epithelial Cell Basal Medium and Growth kit, ATCC, VA, USA) in T75 flasks. Media was changed three times per week. Upon confluency, cells were detached using 0.05% Trypsin-EDTA at 37°C and split at a 1:2 ratio. After 3 passages, the cells were harvested for characterization.

### hiPSC culture and differentiation

Cryopreserved hiPSCs were thawed and plated into 6-well culture plates coated with 1.1 µg/cm^2^ recombinant human laminin-521 (rhLN-521, A29248, Gibco) in 2 ml mTeSR Plus medium (100-0274, STEMCELL). The medium was changed every other day, and the cells were passaged using 0.5 mM EDTA in PBS upon confluence and reseeded on rhLN-521 coated 6-well plates (0.55 µg/cm^2^). For induction of surface ectoderm differentiation, the cells were dissociated using 0.5 mM EDTA for 4–5 min at 37°C and resuspended as single cells after centrifugation at 200 g for 5 min. Then, 5 X 10^5^ iPSCs were transferred into each well of 6-well culture plates coated with rhLN-521 (0.55 µg/cm^2^) in Essential 6 medium (E6, A1516401, Gibco) supplement with 10 mM Y-27632 (ab120129, Abcam) on day 1, 10mM SB-505124 (S4696, Sigma-Aldrich) and 50 nM bFGF (78003, STEMCELL) on day 2, and 25 nM BMP-4 (78211, STEMCELL) on days 3 and 4. During the induction phase, the medium was changed every day and small molecules were added to the media accordingly. From day 5 to 25, the cells were grown in DKSFM (0744019, Gibco) and the medium was changed every other day. The differentiated cells were harvested at days 5, 10, 15 and 25 (D5 to D25) by incubating with 0.05% Trypsin-EDTA at 37°C until most of the cells were round (approximately 8–10 min) and used for quantitative real-time PCR (qPCR) and/or immunocytochemistry (ICC).

### Quantitative real-time PCR

Total RNA was extracted using TRIZOL reagent (ThermoFisher) and cDNA was synthesized using the RT^2^ First Strand Kit (Qiagen). qPCR reactions were prepared using RT^2^ SYBR Green qPCR Mastermixes (Qiagen) and performed using the CFX Connect Real-Time System (BioRad Laboratories, 45 cycles, 95 °C for 20 s, 60 °C for 60 s). Each reaction was repeated 3 times (technical replicates). Gene expression values were normalized against *GAPDH* expression and relative expression compared with hiPSCs were calculated. The mRNA expression levels for the selected markers are presented as mean values. Error bars indicate standard deviation. See Table 1 for a list of primers used for target genes.

**Table 1.**
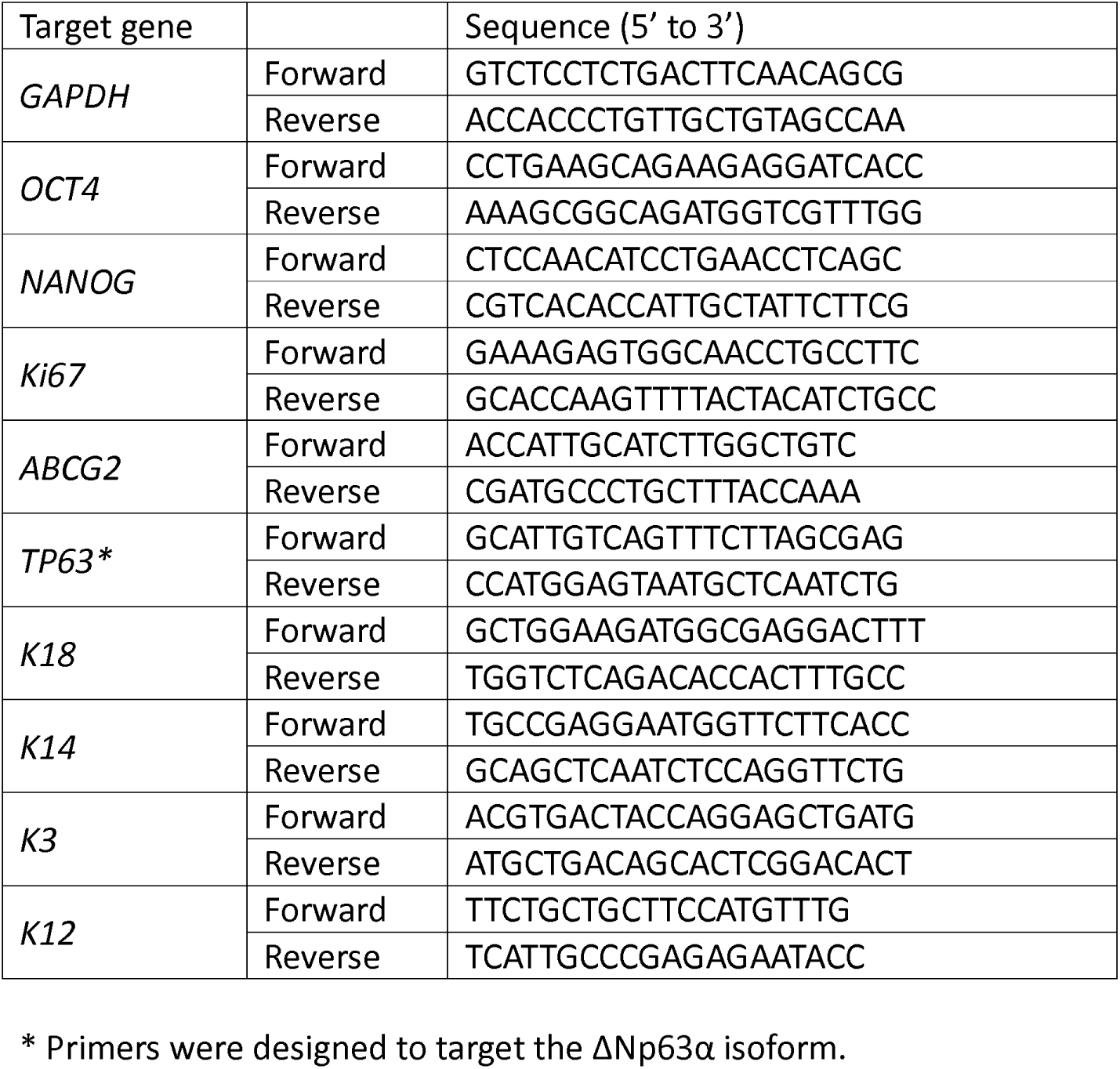
Primers used for quantitative real-time PCR.

### Immunocytochemistry

ICC was performed on D10 and D15 post differentiation and HCEC. The cells were washed with 1X Dulbecco’s Phosphate-Buffered Saline (DPBS) three times and then incubated with 4% paraformaldehyde for 20 min at 37°C and then permeabilized with 0.1% Triton-X100 for 15 min at room temperature (RT). The cells were washed again three times with 1X DPBS and incubated with 2% bovine serum albumin (BSA) at RT for 1h. Primary antibodies (diluted in 0.2% BSA) were added and incubated at 4°C overnight. After washing three times with 1X DPBS, secondary antibodies and nuclear counterstain (DAPI, Sigma-Aldrich) were added and incubated at RT in dark for 45 min (see Table 2 for details of primary and secondary antibodies and concentrations). The wells were then washed three times with PBS-T (DPBS with 0.1% TWEEN 20), air dried, and kept in 1X DPBS until imaging. Fluorescent imaging was performed using Nikon Eclipse N*i* microscope equipped with DS-*Qi2* camera and analyzed using NIS-Elements Viewer (version 5.41.00; Laboratory Imaging) and ImageJ software (NIH). Quantitative measurement of the total cell number and ratio of cells positive for p63α, ΔNp63α, and Ki67/ΔNp63α double positive were performed on D10 and D15. For this purpose, multiple images were captured with 10X magnification and at least 5 images (1.2 mm x 1.2 mm) and at least 500 cells were analyzed. The total number of cells were measured by counting the nuclei stained with DAPI and the ratio of positive cells were calculated by dividing the number of cells positive for the target antibody target by the number of DAPI-positive cells.

**Table 2.**
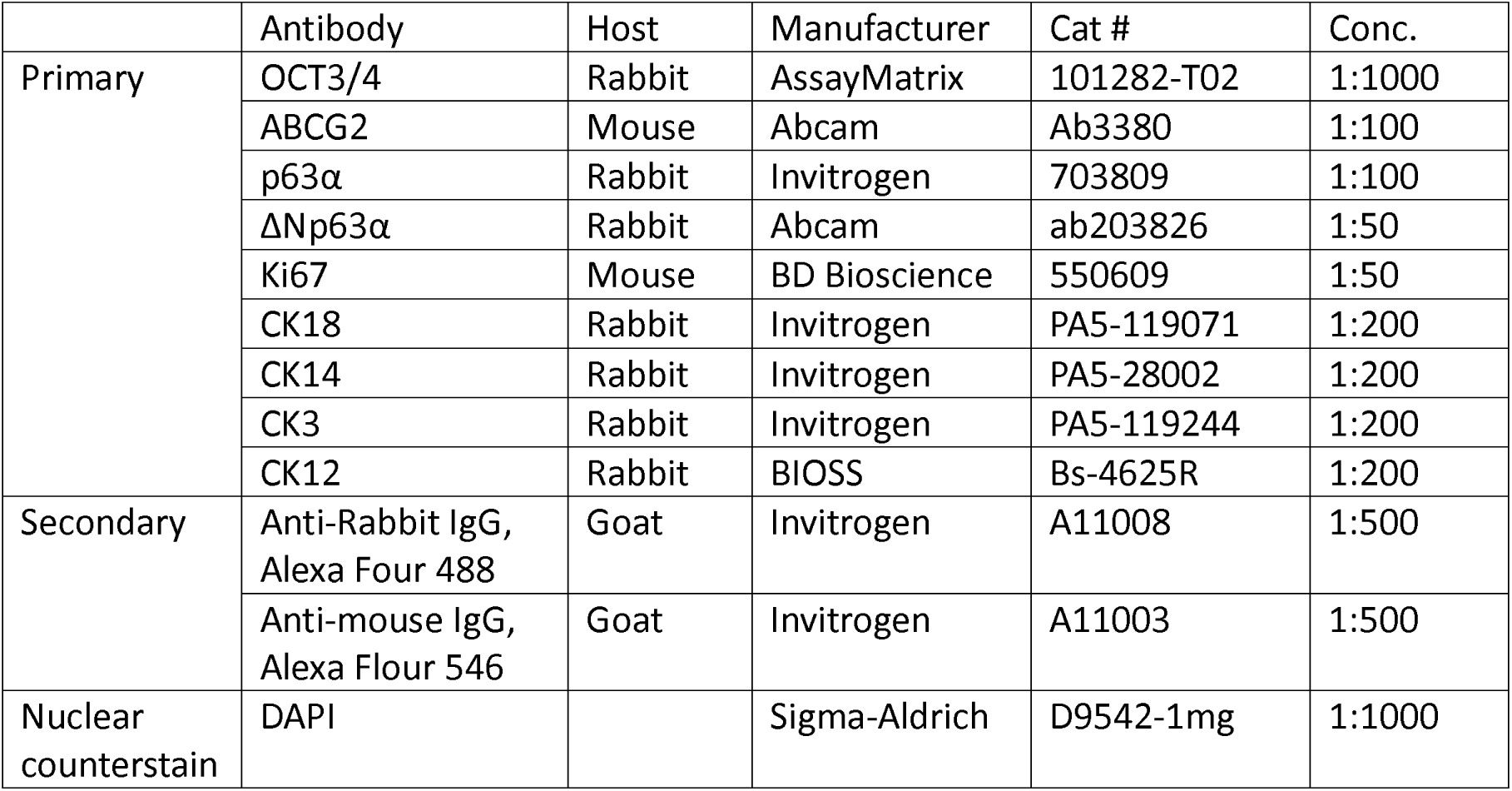
Primary and secondary antibodies used for immunocytochemistry.

### Statistical analyses

Gene expression and cell counting data were recorded in the statistical package for social sciences (SPSS) version 23 (SPSS/IBM. Inc., Chicago, IL, USA). Normality of data was tested using the Shapiro-Wilk Test and appropriate tests were applied to compare relative expressions, total number of cells and ratio of positive cells between different cell lines and post differentiation days. Independent samples t-test and Mann-Whitney U test were used for parametric and nonparametric variables, respectively. P-values less than 0.05 were considered significant.

## Results

Within 5 days following induction, cell morphology started to change toward large epithelial-like cells with a polygonal shape mixed with undifferentiated (UD) colonies (Figure 1). From D10 to D25, epithelial-like cells became predominant and most of the cells were polygonal shape on D25 (Figure 1). The cell density was 324 cells/mm^2^ in D10 and 149 cells/mm^2^ in D15 (54% decline, P=0.21).

**Figure 1.**
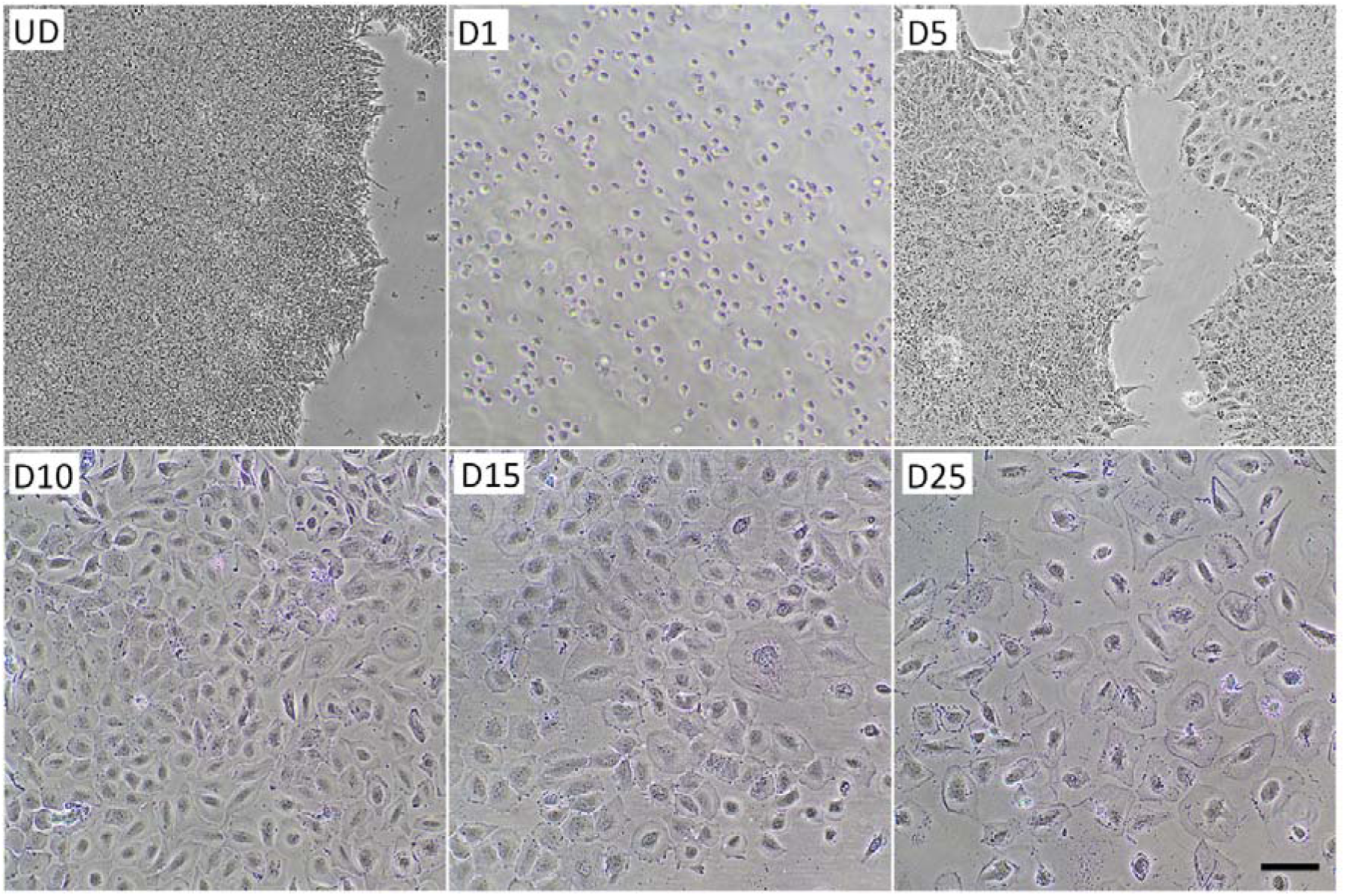
Bright-field microscopy shows undifferentiated hiPSCs (UD) and 1 day after initiation of ectodermal induction in single cells (D1). Epithelial-like cells started to appear on day 5 (D5). On day 10 (D10), there was a monolayer of predominant epithelial-like cells. The cell density declined on day 15 (D15), with an apparent senescence on day 25 (D25). Scale bar = 100 microns.

### Pluripotency and proliferation markers

Following ectodermal induction (D5), the expression of *OCT4* and *NANOG* declined significantly compared with UD cells. Although the expression of both markers increased significantly on D10 compared with D5, they were still significantly lower than UD cells. On D25, the expression of *NANOG* remained significantly lower than UD but higher than D5. D25 cells expressed a significantly higher level of *OCT4* compared with UD, D5 and D10. The HCEC line expressed significantly lower *OCT4* and *NANOG* mRNAs compared with UD cells and all differentiation time points (Figure 2). Expression of *Ki67* mRNA was significantly higher in UD and D10 cells compared with D5, D25, and HCEC (Figure 2).

**Figure 2.**
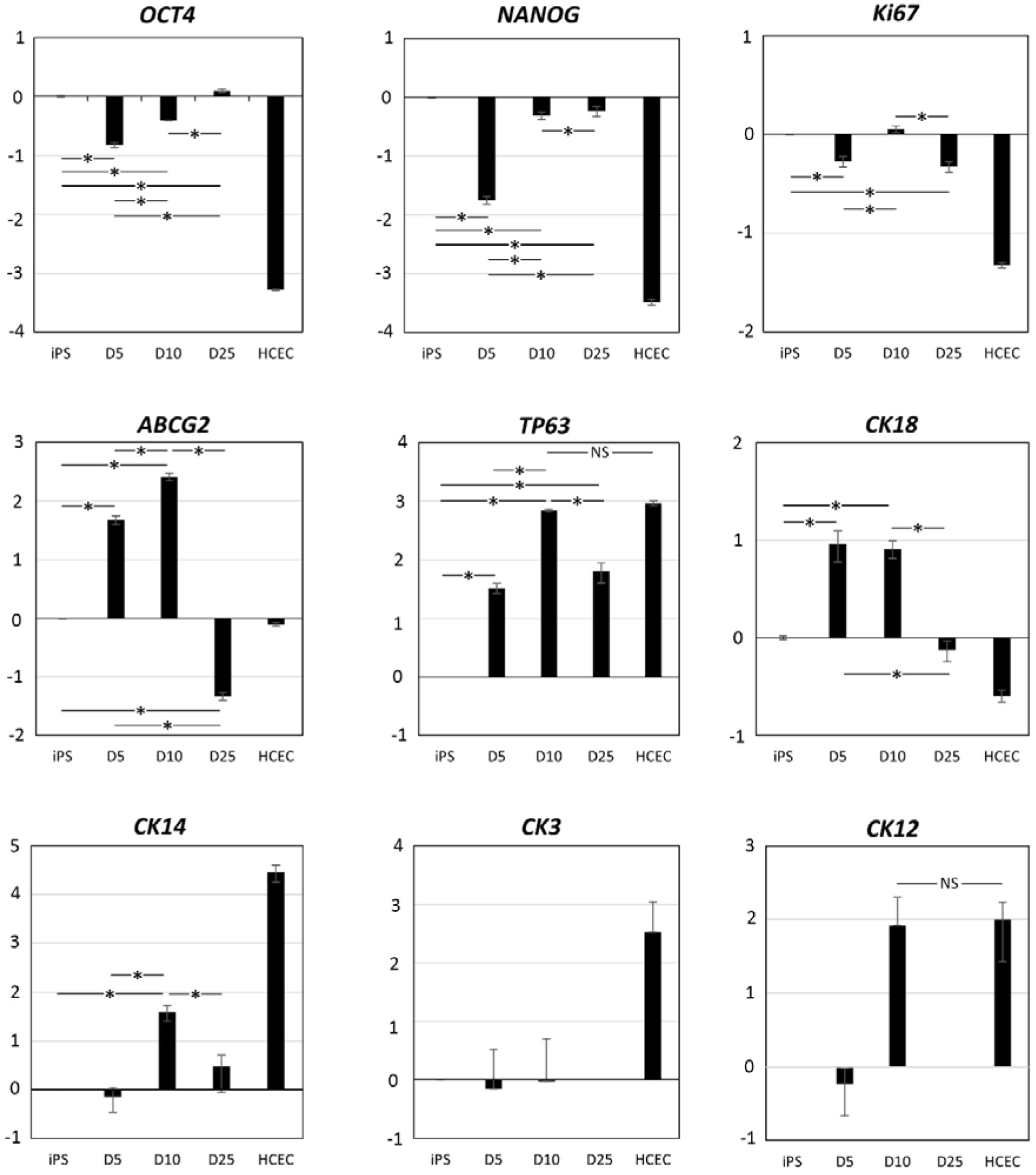
Quantitative real-time PCR shows a significant reduction in the expression of pluripotency (*OCT4* and *NANOG*) and proliferation (*Ki67*) markers immediately after the ectodermal commitment (i.e., day 5). In contrast, limbal stem cell markers (*ABCG2* and *TP63*) and early epithelial differentiation marker (*CK18*) were overexpressed significantly. Peak expression levels of *ABCG2*, *TP63*, *CK14*, and *CK12* mRNAs were observed on day 10. Expression levels are normalised to *GAPDH*. Significant differences and important non-significant differences are marked. Note that the expression levels of HCEC were significantly different with iPSC and all time points of differentiation for all markers, except *TP63* and *CK12* on day 10. *CK3* and *CK12* mRNAs were not detected on day 25. *P < 0.05. HCEC = human corneal epithelial cell; NS = not significant.

### Limbal stem cell markers

Expression of *ABCG2* mRNA was significantly higher in D5 and D10 compared with UD cells, but significantly lower than UD cells in D25 cells and HCECs. Also, D10 cells expressed significantly higher levels of *ABCG2* mRNA compared with D5, D25, and HCECs (Figure 2). Expression of *TP63* mRNA was significantly higher in D5, D10, and D25 cells and HCECs compared with UD cells. In addition, D10 and HCECs expressed higher levels of *TP63* compared with D5 and D25, while there was no significant difference between D10 and HCEC (Figure 2). ICC showed weak staining for ABCG2 and strong nuclear staining in a proportion of D10 cells (Figure 3). The percentage of p63α-positive cells was 71% and 56% in D10 and D15, respectively (P=0.022). The percentage of ΔNp63α-positive cells was 70% and 51% in D10 and D15, respectively (P=0.049). Double staining for ΔNp63α and Ki67 showed that 57% and 41% of cells were positive for both markers in D10 and D15, respectively (P=0.056) (Figures 4 and 5).

**Figure 3.**
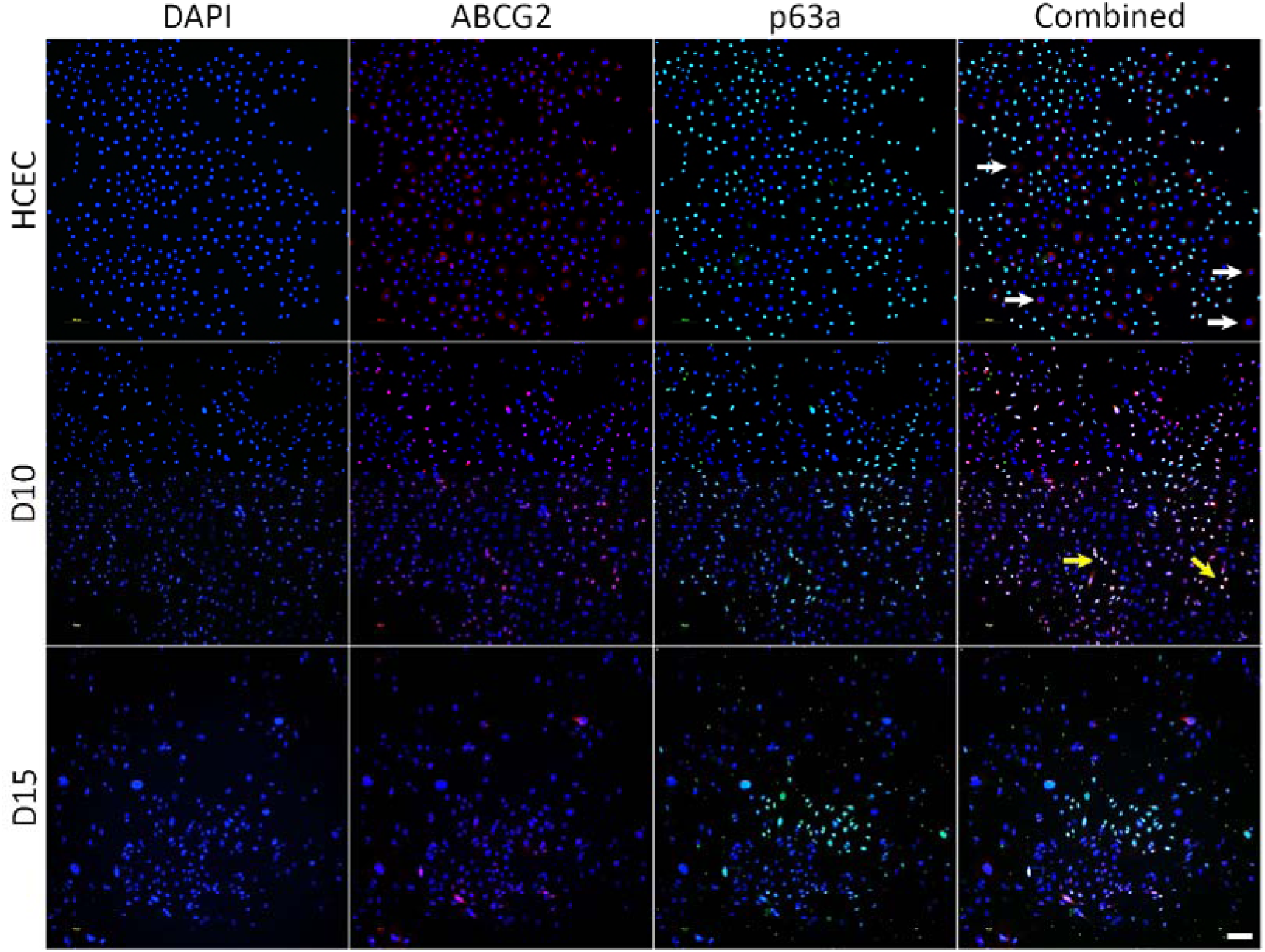
Immunocytochemistry for limbal epithelial progenitor markers shows positive staining for ABCG2 and p63α in the human corneal epithelial cell (HCEC) line and D10 and D15 differentiated iPSCs. Interestingly, ABCG2-positive/p63α-negative cells (white arrows) were observed in the HCECs, but not in the differentiated D10 and D15 cells. In addition, small populations of double-positive cells were observed in D10 cells (yellow arrows). Scale bar = 100 microns.

**Figure 4.**
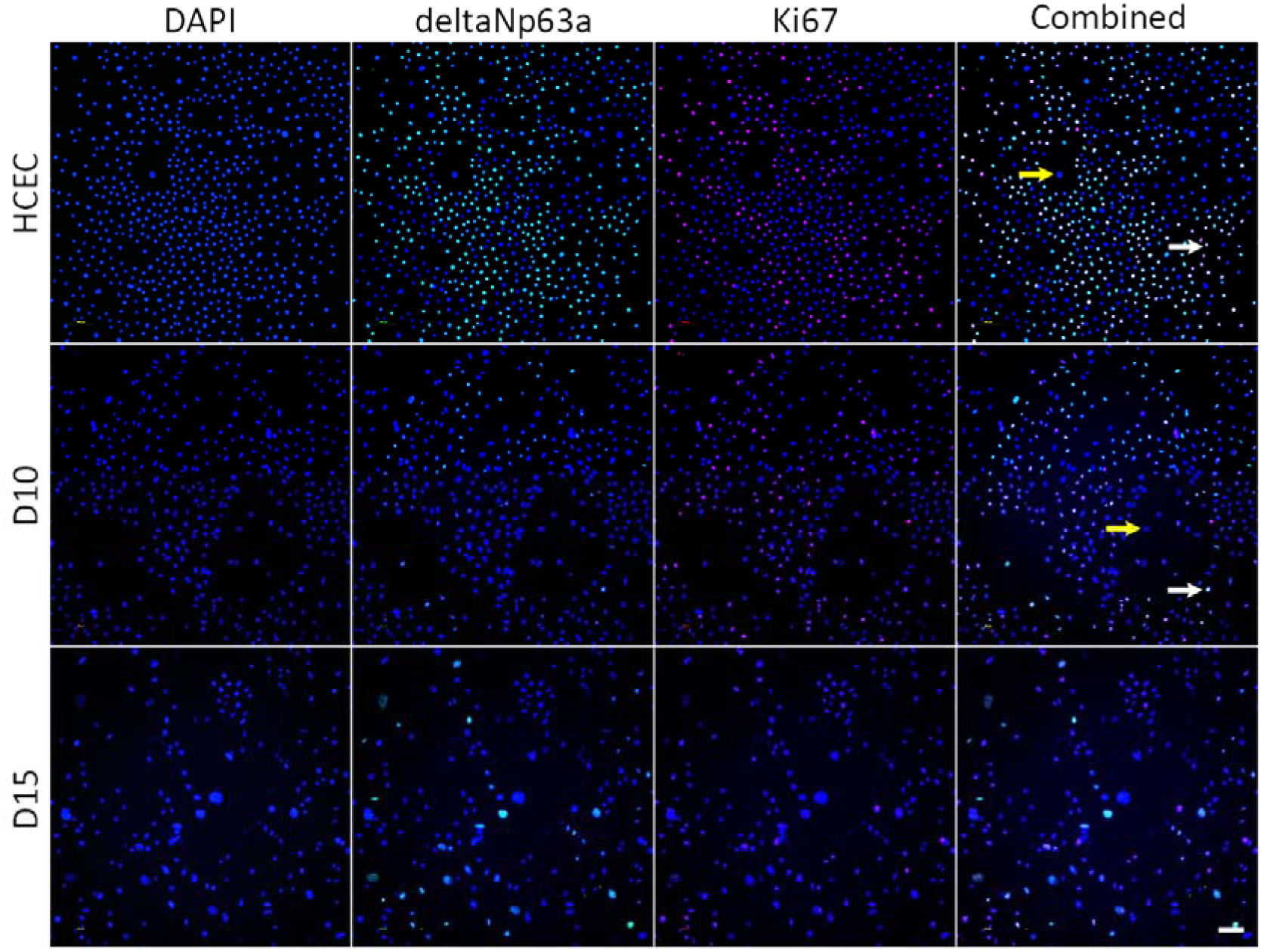
Double staining for limbal epithelial marker ΔNp63α and proliferation marker Ki67 showed abundant double-positive cells in both HCEC and D10 (white arrows). There were also double-negative cells (yellow arrows) which appeared to be terminally differentiated cells. Scale bar = 100 microns.

**Figure 5.**
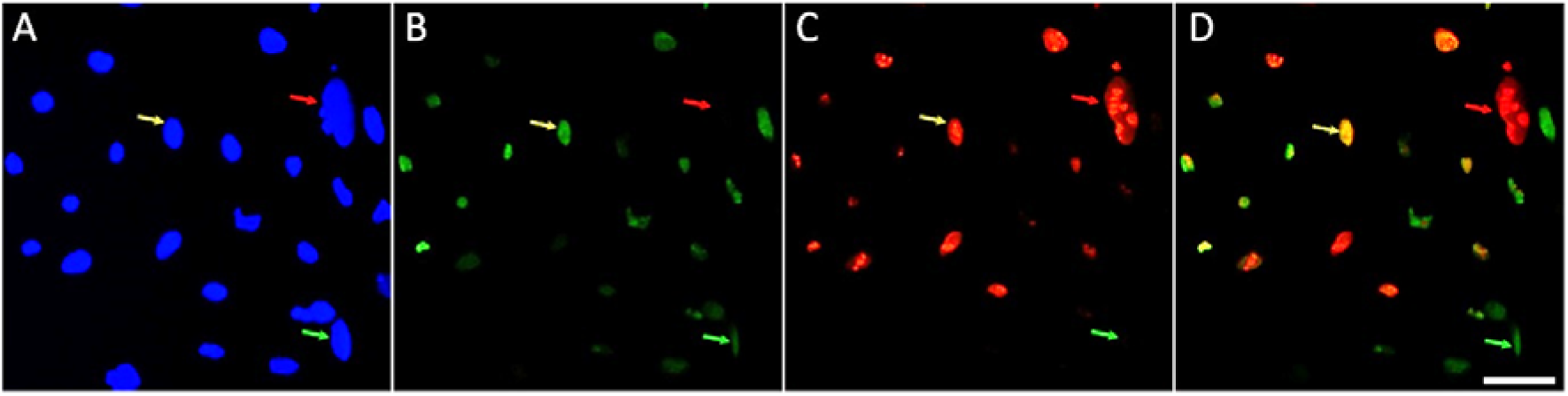
A magnified image of DAPI (A), ΔNp63α (B), Ki67 (C) and combined (D) staining in D10 shows cells only positive for ΔNp63α (green arrows), only positive for Ki67 (red arrows), or positive for both markers (yellow markers). Scale bar = 50 microns.

### Simple and stratified squamous epithelial markers

Ectodermal induction resulted in an approximately 10-fold increase in *CK18* mRNA level in D5 and D10 compared with UD cells (P<0.05). D25 and HCEC expressed significantly lower levels of *CK18* mRNA compared with D5 and D10 (Figure 2). *CK14* mRNA was significantly higher than UD, D5, and D25 in D10 (P<0.05). However, it was still significantly lower than HCEC (Figure 2). Both CK18 and CK14 proteins were produced in differentiated cells in D10 and D15. Whilst staining for CK18 in D10 and D15 was stronger than HCEC, CK14 staining was stronger in HCECs compared with D10 and D15 cells (Figure 6).

**Figure 6.**
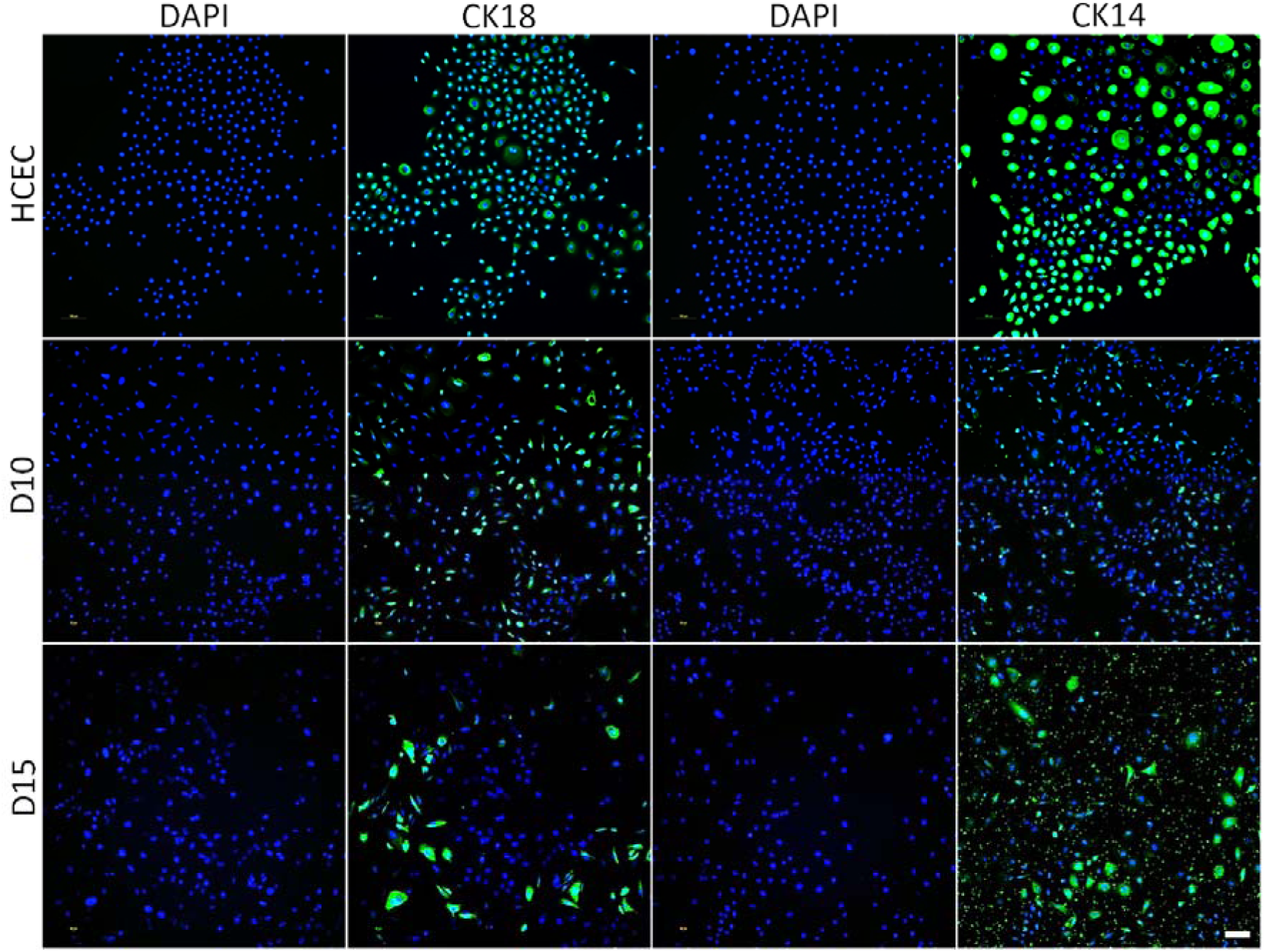
Populations of CK18-positive and CK14-positive cells appeared on day 10 post-differentiation and were characterized by colonies of large CK18 and CK14-positive cells in D15. The staining patterns in HCECs are shown as an example. Scale bar = 100 microns.

### Corneal epithelial markers

*CK3* mRNA expression did not change significantly in the differentiated cells compared with UD cells and it was not detected in D25 (Figure 2). *CK12* mRNA was significantly higher in D10 compared with UD and D5 (P<0.05) and it was not detected in D25. The level of *CK12* mRNA in D10 and HCEC was approximately 100-folds higher than the UD cells (P<0.05), but it was not significantly different between D10 and HCEC (Figure 2). D10 cells stained positive for CK3 and CK12. Large epithelial cells strongly positive for CK3 appeared on D15 (Figure 7).

**Figure 7.**
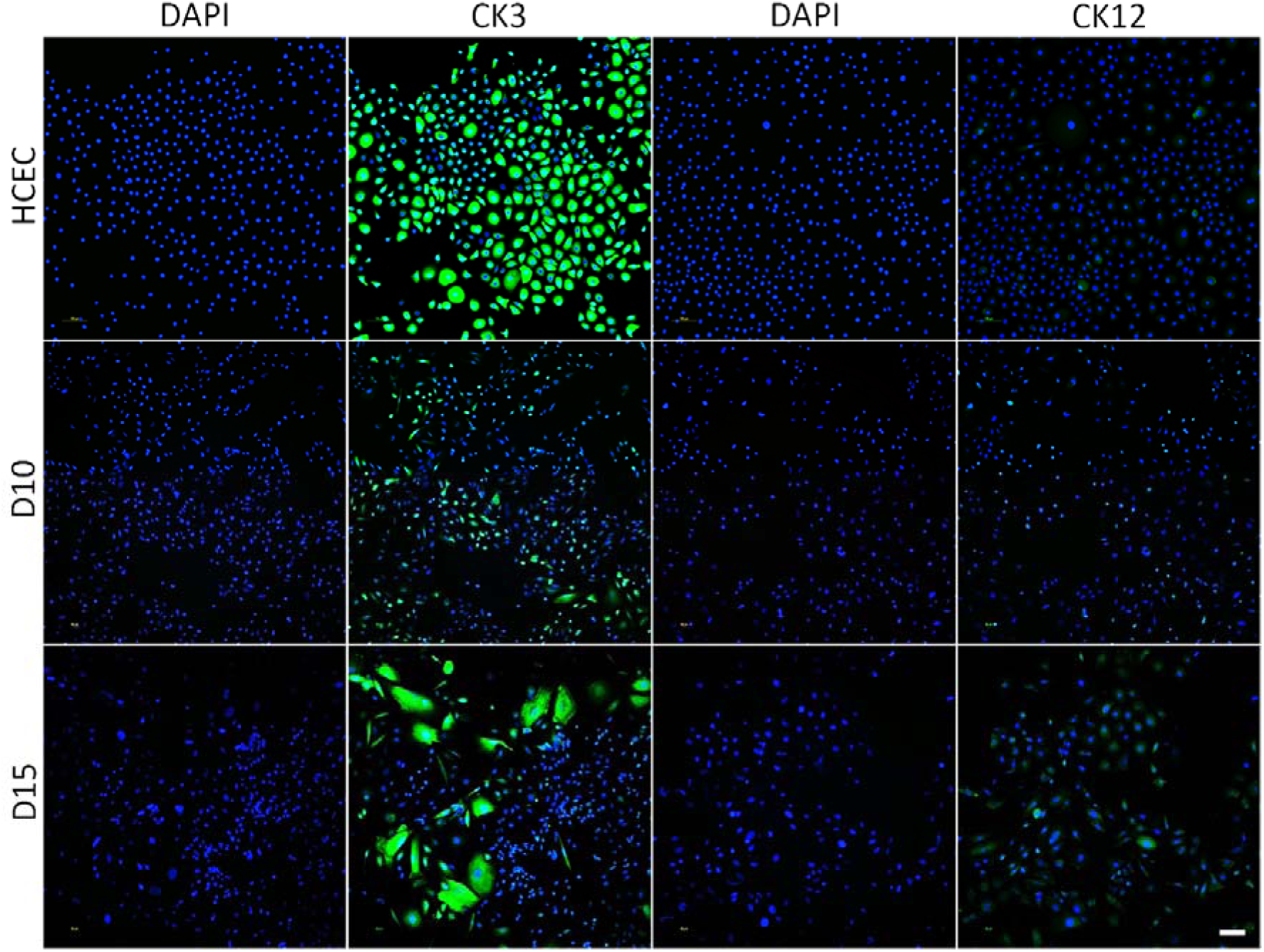
CK3 and CK12-positive cells were observed in both D10 and D15 cells. Mature epithelial morphology and strong CK3 staining might be an indicator of terminal differentiation of D15 cells, which may explain their senescence in D25. The staining patterns in HCECs are shown as an example. Scale bar = 100 microns.

## Discussion

Various modifications of xeno-free and serum-free methods have been described for the differentiation of human pluripotent stem cells (hPSCs), i.e., human embryonic stem cells (hESCs) and hiPSCs, into LESCs.^17,18^ Current methods incorporate a two-stage protocol including an ectodermal induction followed by an epithelial differentiation phase. There is robust evidence that inhibition of TGF-β and activation of bFGF and BMP-4 pathways in the early stages of differentiation promote limbal epithelial differentiation of hPSCs.^19,20^ We used an adherent culture system and E6 medium supplemented with a TGF-β inhibitor, bFGF, and BMP-4 in the induction phase, which resulted in a significant downregulation of pluripotency markers similar to previous studies that used embryonic body (EB) formation in suspension culture system for the ectodermal induction.^12,21,22^ Yang *et al.* also used adherent culture for induction in E6 medium and reported significant downregulation of the pluripotency markers after 2 weeks.^17^ Ectodermal induction using a suspension culture in E6 supplemented with SB-505124, IWP-2, and bFGF resulted in a high expression of LESC markers and low expression of pluripotency markers after 14 days.^22^ These findings suggest that the optimal time for starting differentiation depends on the induction and differentiation methods.

We observed a significant increase in *OCT4* and *NANOG* expression during the differentiation phase (i.e., from D5 to D25). Previous studies have shown that human limbal epithelial cells can express *OCT4* and *NANOG* mRNA^23^ and hPSC-derived LESCs express high amounts of pluripotency marker mRNA and protein after 21 days.^24,25^ We also observed a significant increase in the expression of *TP63* and *ABCG2* in parallel to the downregulation of the pluripotency markers following the induction phase. Despite the upregulation of *TP63*, which has been a constant finding in the literature, the findings regarding *ABCG2* are controversial. It was shown that human limbal epithelial cells express higher levels of *ABCG2* mRNA compared with central corneal epithelium.^23^ A study showed that while hPSC-derived LESCs had higher *ABCG2* and lower *TP63* expression levels after 10–11 days of differentiation, day 24–25 cells expressed low levels of *ABCG2* and high levels of *TP63* mRNAs.^26^ In contrast, other studies have shown no^17^ or minimal^25^ increase in the expression of *ACBCG2* during the induction and differentiation phases, despite consistent increase in *TP63* and other LESC markers.

We used DKSFM for the differentiation phase, which is a serum-free medium optimized for the isolation and expansion of human keratinocytes without the need for bovine pituitary extract (BPE) supplementation or the use of fibroblast feeder layers. Our method resulted in 70% ΔNp63α-positive cells, of which 80% were also positive for Ki67, after 10 days of initiation of induction. Hongisto and colleagues used similar small molecules for the induction and CnT-30 for differentiation and achieved 66% ΔNp63α-positive cells after 3.5 weeks. Interestingly, the ratio of ΔNp63α-positive cells increased to 80–90 % after recovery from cryostorage.^21^ Yang and colleagues used E6 for the induction and differentiation of hESCs into LESCs and reported 92% p63α-positive cells after 10 days, which was higher than our study. They also reported that by day 50, nearly 50% of cells positive for CK15 were also positive for Ki67. They concluded that E6 alone (supplemented with small molecules during the induction phase) is enough for differentiating hPSCs into limbal and corneal epithelial cells.^17^

While we achieved a high ratio of ΔNp63α/Ki67-positive cells in less than 2 weeks, CnT-30 and E6 resulted in high proportion of LESC-like cells over 3–4 weeks and 6–8 weeks, respectively. However, we observed a significant decrease in the total number of cells as well as the ratio of p63α and ΔNp63α-positive cells at D15 compared with D10. Interestingly, although the percentage of ΔNp63α/Ki67-positive cells reduced from 57% in D10 to 41% in D15, the ratio of ΔNp63α-positive cells that stained positive for Ki67 remained stable (80% in D10 and D15). At the gene expression level, D25 cells revealed significantly lower levels of *Ki67*, *ABCG2*, *CK15*, and *CK18*, compared with D10. Mikhailova and colleagues reported that the percentage of p63α-positive cells reduced from 95% on day 30 to 70% on day 44 post differentiation.^12^ One explanation for this phenomenon could be that prolonged culture in the differentiation medium may induce terminal differentiation of the progenitor cells, resulting in the gradual reduction in the proliferation potential of the differentiating cells. According to these findings, this process might require longer time in E6 and CnT-30 media, compared with DKSFM.

Our results were compatible with the embryonic stages of limbal epithelial development as shown by an early increase in *CK18* expression on D5 followed by upregulation of *CK14* on D10, which was also observed in previous studies. For instance, Shalom and colleagues have shown that following induction in a conditioned medium supplemented with BMP-4, the peak levels of *CK18* and *CK14* were observed after 4 and 8 days, respectively.^11^ Similarly, Lian and colleagues reported that CK18 was the first marker that increased following ectodermal induction and that further differentiation of CK18-positive cells in DKSFM resulted in a high percentage (90%) of CK14-positive cells after two weeks.^18^

The main limitation of our study is using a single hiPSC line. Also, using flow cytometry could be useful in quantification of the cells positive for limbal-specific markers. In addition, a more comprehensive characterization using a broader range of markers could be useful in further characterization of the differentiated cells and determining the nature of the other cells.

## Conclusion

Our study showed that ectodermal induction in E6 supplemented with known small molecules in an adherent culture system followed by epithelial differentiation in DKSFM results in differentiation of hiPSCs into LESC-like cells within 2 weeks. However, these results should be validated in multiple biological replicates and by more comprehensive characterization. In addition, changing the medium from differentiation medium to growth medium (i.e., a three-stage protocol) might be useful for promoting expansion of the differentiated cells instead of terminal differentiation of the LESC-like cells.

